# Transposable elements drive the evolution of metazoan zinc finger genes

**DOI:** 10.1101/2022.11.29.518450

**Authors:** Jonathan N. Wells, Ni-Chen Chang, John McCormick, Caitlyn Coleman, Nathalie Ramos, Bozhou Jin, Cédric Feschotte

**Author notes:** These authors contributed equally to this work (Ni-Chen Chang, John McCormick).

## Abstract

Cys2-His2 Zinc finger genes (ZNFs) form the largest family of transcription factors in metazoans. ZNF evolution is highly dynamic and characterized by the rapid expansion and contraction of numerous subfamilies across the animal phylogeny. The forces and mechanisms underlying rapid ZNF evolution remain poorly understood, but there is growing evidence that the targeting and repression of lineage-specific transposable elements (TEs) plays a major role in the diversification of the Kruppel-associated box ZNF (KZNF) subfamily, which predominates in tetrapod genomes. At present, it is unknown whether this function and co-evolutionary relationship is unique to KZNFs, or a broader feature of metazoan ZNFs. Here, we present evidence that genomic conflict with TEs has been a central driver in the diversification of ZNFs in animals. Sampling from more than 4000 animal genome assemblies, we show that the copy number of retroelements correlates with that of ZNFs across at least 750 million years of metazoan evolution, both within and between major taxonomic groups. Using computational predictions, we show that ZNFs preferentially bind TEs in a diverse set of representative animal species. We further investigate one of the most expansive ZNF subfamilies found in cyprinid fish, which are characterized by a conserved domain we dubbed the **Fi**sh **N**-terminal **Z**inc-finger associated (FiNZ) domain. FiNZ-ZNFs have dramatically expanded in several fish species, including the zebrafish in which we predict ~700 FiNZ-ZNF genes. Almost all are located on the long arm of chromosome 4, and recent duplicates are evolving adaptively under positive selection. Like mammalian KZNFs, the bulk of zebrafish FiNZ-ZNFs are expressed in waves at the onset of zygotic genome activation. Blocking FiNZ-ZNF translation using morpholinos during early zebrafish embryogenesis results in a global de-repression of young, transcriptionally active TEs, likely driven by the failure to establish heterochromatin over these elements. Together, these data suggest that ZNF diversification has been intimately connected to TE expansion throughout animal evolution and that families of ZNFs have been deployed independently in fish and mammals to repress TEs during early embryogenesis.

## Introduction

The Cys2-His2 Zinc Finger (ZNF) is a short nucleic acid binding domain widespread in eukaryotes. ZNFs are often arranged in tandemly repeated arrays, resulting in proteins with versatile nucleic acid binding ability (Wolfe et al., 2000; Najafabadi et al., 2017; Klug, 2010). In metazoans tandem-repeat ZNF genes have undergone dramatic lineage-specific expansions in gene copy number, as well as an increase in the diversity of DNA sequences that they recognize (Heger et al., 2020; Najafabadi et al., 2017). Many ZNF proteins contain accessory domains that modulate chromatin states and transcription activity, such as the repressive **Kr**üppel-**a**ssociated **b**ox (KRAB) (Bellefroid et al., 1991). Thus, ZNFs represent the largest and most dynamically evolving transcription factor family in metazoans.

The abundance and diversity of C2H2 ZNF genes is a hallmark feature of metazoan genomes (Najafabadi et al., 2017; Heger et al., 2020), and in species whose genomes have been sequenced, ZNF gene copy number spans three orders of magnitude, from fewer than ten in many nematodes, to hundreds, if not thousands in vertebrates, arthropods, and mollusks (Albertin et al., 2015; Goodstadt et al., 2007; Panfilio et al., 2019). Their lineage-specificity can be seen through comparisons of closely related species. For example, a survey of three great ape species and rhesus macaque found that approximately 10% (213 out of 2,253) of predicted KRAB-ZNFs (KZNFs) were species-specific, including 7 from humans and 145 in orangutangs (Nowick et al., 2011). This rapid evolution contrasts with most other transcription factors, which are deeply conserved across metazoans (Degnan et al., 2009).

However, the same qualities that differentiate ZNFs from other transcription factors make them challenging to study, and consequently there have been few attempts to produce a unified theory explaining their prevalence and recurrent expansions (Emerson and Thomas, 2009; Fedotova et al., 2017). Early hints at the function of the KZNFs came from work showing that deletion of the genes *TRIM28* and the K3K9 methyltransferase *SETDB1* led to dramatic increases in endogenous retrovirus expression in mice (Rowe et al., 2010; Matsui et al., 2010). Since TRIM28 was known to be an important cofactor of the KRAB domain, this led to the proposal that KZNFs are in a coevolutionary relationship with retroelements (Thomas and Schneider, 2011). In this study, we aim to test whether genetic conflict with transposable elements (TEs) is the central force driving the rapid evolution of metazoan ZNFs more generally.

TEs are selfish genetic elements that replicate independently within their host genomes, and in most eukaryotes comprise between 5% and 85% of the genome. While individual TE insertions are most likely to be neutral or deleterious, in aggregate, TEs represent an important source of genetic variation and innovation. For example, in vertebrates TEs are a major source of cis-regulatory elements and coding sequences coopted for host gene regulation (Senft and Macfarlan, 2021; Fueyo et al., 2022; Almeida et al., 2022). On the other hand, uncontrolled TE proliferation has deleterious consequences ranging from insertional mutagenesis and genomic instability caused by ectopic recombination between TE insertions, to dysregulation of gene expression stemming from the fact that many TEs carry their own promoters and regulatory sequences. Consequently, metazoans have evolved a variety of defenses to control their spread, most notably the piRNA system (Aravin et al., 2007; Czech and Hannon, 2016) and the previously mentioned KZNFs (Ecco et al., 2017; Bruno et al., 2019). One hallmark of both of these defense systems is their ability to adapt to changing population of invaders, which leads to selection pressure on the invaders to evade these defenses, thereby creating arms race dynamics (McLaughlin and Malik, 2017).

There is growing evidence that KZNFs are engaged in such arms races with TEs (Bruno et al., 2019; Fernandes et al., 2018; Jacobs et al., 2014). KZNFs form the largest ZNF gene subfamily found in Sarcopterygii, i.e. tetrapods and lobe-finned fish (Imbeault et al., 2017; Bellefroid et al., 1991). The extensive variation in KZNF copy number across species has long been appreciated (Bellefroid et al., 1991; Huntley et al., 2006), but the evolutionary forces driving this remained elusive until a breakthrough study showing that the number of ZNF domains in a given genome is positively correlated to retroelement copy number across a small but diverse sample of vertebrates, suggesting a coevolutionary relationship between the two (Thomas and Schneider, 2011). This relationship was later bolstered by ChIP-seq experiments in humans and mice mapping the genome-wide binding of hundreds of KZNFs, which revealed that most target specific TE families (Imbeault et al., 2017; Wolf et al., 2020). Furthermore, KZNF knockouts in mouse and humans lead to upregulation of TE expression (Haring et al., 2021; Wolf et al., 2020). Mechanistic studies showed that TE transcriptional repression via KZNF proteins is typically mediated via their KRAB domain, which interacts with KAP1/TRIM28 corepressor to recruit the H3K9me3 writer, SETDB1, amongst several other chromatin silencing factors (Wolf and Goff, 2009; Rowe et al., 2010; Matsui et al., 2010; Ecco et al., 2017); in doing so, most KZNFs nucleate the formation of heterochromatin at their target TE loci. Together, these findings support the idea that TE proliferation is a driving force behind the diversification of KZNFs.

Much less is known about the factors driving the evolution of other ZNF families. KZNFs represent a small fraction of all ZNFs, being restricted to some thirty thousand tetrapods, against a background of millions of animal species (Mora et al., 2011; Sahney et al., 2010), most of which are predicted to harbor hundreds of individual ZNF genes. One of the few other large families to have been studied are the zinc finger-associated domain (ZAD)-ZNFs, which are found in many insect species (Chung et al., 2007). The handful of ZAD-ZNFs characterized in *Drosophila melanogaster* do not appear to have clear roles in TE repression, but instead perform a variety of functions related to heterochromatin organization (Kasinathan et al., 2020). Thus far, only one ZAD-ZNF gene (CG17801) has been reported to directly affect TE transcript levels, and was identified as part of an earlier screen for piRNA pathway components (Czech et al., 2013). More recently, the ZAD-ZNF protein Kipferl has been shown to regulate TE activity indirectly, by targeting the protein Rhino to heterochromatic piRNA clusters in *Drosophila melanogaster*, thus licensing piRNA production from these loci (Baumgartner et al., 2022). Despite apparent differences in the immediate function of most ZAD- and KZNF genes, both families share features that are broadly characteristic of the majority of ZNF genes thus far studied, namely: rapid turnover and sequence evolution, evidence of being under positive selection, involvement in establishing or maintaining heterochromatin, and expression during early embryogenesis (Chung et al., 2007; Kasinathan et al., 2020; Ecco et al., 2017). Other highly expanded ZNF families, such as those of hemipteran bugs, octopus, and zebrafish, also possess subsets of these features (Panfilio et al., 2019; Albertin et al., 2015; Howe et al., 2016).

Here, we investigate the hypothesis that interaction with TEs is a driving force underlying the molecular evolution and functional diversification of ZNF families across metazoans. Through a survey of all currently available metazoan genome assemblies, we find that the number of ZNF open reading frames is positively correlated with the number of retroelements across a large and diverse sample of animals. Using in silico predictions of ZNF binding specificity, we show that ZNFs from a sample of distantly related animals preferentially recognize TEs, as has been shown for KZNFs. We then turn our focus to a large family of ZNFs present in zebrafish, which are defined by its association with a conserved protein domain we have termed **Fi**sh **N**-terminal **Z**inc-finger associated (FiNZ). These genes are unique to cyprinid fish and have evolved in parallel to the KZNFs found in tetrapods. Using refined gene annotations, we find that in zebrafish, recently duplicated FZNF paralogs are under positive selection, consistent with ongoing arms races with TEs. Finally, by simultaneously targeting over 400 FZNF genes for knockdown during zebrafish embryogenesis, we show that FZNF depletion causes de-repression of TEs.

## Results

### Annotation of ZNFs and TEs in metazoan genomes

We sought to estimate the number of ZNF genes in a large set of publicly available metazoan genome assemblies. ZNF genes are challenging to annotate due to their highly repetitive nature and restricted patterns of mRNA expression. Consequently, even in well curated genome assemblies, the true number of functional ZNF genes is likely underestimated (Huntley et al., 2006; White et al., 2017). To avoid biases caused by incomplete gene annotation, we used profile-HMM searches with HMMER (Eddy, 2011) to identify ZNF open reading frames (ORFs), here defined as any stretch of protein coding sequence, regardless of whether they are initiated by a start codon. As our aim in this project was to investigate tandem-repeat ZNF genes, we restricted our search to ORFs with at least five ZNF domains. Searching 3,221 metazoan genome assemblies revealed extensive variation across species, from fewer than ten ZNF ORFs in most nematode worms, to upwards of a thousand in many vertebrates, mollusks and arthropods (Fig 1A, Supp. Data 1-2). Since estimates of ZNFs could be biased by genome assembly quality, we set a minimum scaffold N50 of 50 kb to filter out low quality assemblies. To ensure that this threshold was appropriate we calculated the correlation between our ZNF ORFs counts and scaffold N50 length (Supp. Fig. 1A). This revealed a significant, but weak correlation of rho = 0.1, reassuring us that assembly quality was not unduly biasing our ORF counts.

**Figure 1.**
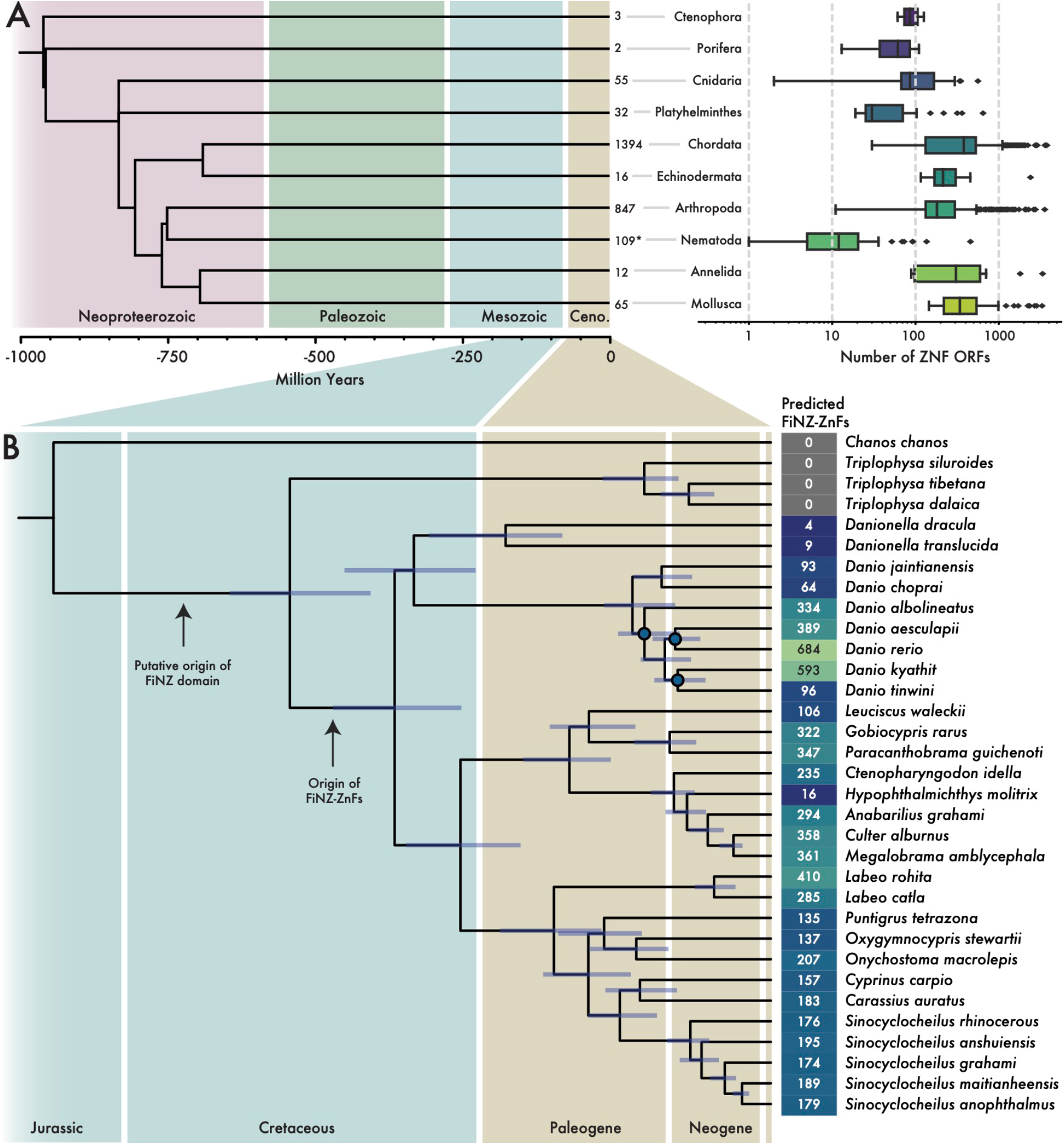
Annotation of ZNF ORFs and full genes in metazoan genome assemblies. A) Boxplots showing distributions of ZNF ORF counts per species in each major taxonomic phylum. Adjacent to boxplots are the number of representative species per phylum; *Number by nematodes excludes those with zero tandem repeat ZNF ORFs (defined as containing at least 5 repeats). Phylogeny acquired from TimeTree5 (Kumar et al., 2022). B) Maximum likelihood phylogenetic tree of Cypriniformes species with high quality genome assemblies. Numbers represent counts of FZNF genes using improved gene annotations. Nodes marked in blue have lower than 95/95 support for Shimodaira-Hasegawa approximate likelihood ratio test/ultra-fast bootstraps, respectively.

In addition to automated counts, we carried out more thorough gene annotations in cyprinid fish (Supp. Data 2). Cyprinids, which include the widely used model organism *Danio rerio* along with many economically and culturally important carp species, are one of the most species-rich vertebrate families, with over 1,500 extant members (Fricke et al., 2020). They possess a large family of ZNF genes characterized by the presence of a 28-amino acid N-terminal domain we dubbed the FiNZ domain (Supp. Fig. 2A). The domain has no significant similarity to known protein domains but is highly conserved across cyprinid ZNFs and enriched in negatively charged glutamic acid residues (p = 3e-06, relative to sequences in the SwissProt51 database (Vacic et al., 2007)). The canonical structure of FinZ-ZNF (FZNF) genes resembles that of KZNFs, with the FiNZ domain being encoded by a single exon upstream of a second exon consisting of an array of ZNF domains (Imbeault et al., 2017). However, the number of copies of the FiNZ domain contained in the first coding exon varies between species – for example, in zebrafish the FiNZ exon almost always contains a single, stand-alone copy, whereas in goldfish it contains up to four tandemly repeated copies.

Using an iterative approach, we annotated putative FZNF genes using a combination of blast (Altschul et al., 1990), HMMER (Eddy, 2011) and Augustus-PPX (Hoff and Stanke, 2018; Keller et al., 2011). In zebrafish, this approach yielded a total of 684 FZNF genes, a substantial increase from the number annotated in the preexisting RefSeq and Ensembl gene sets (Supp. Fig. 2B). Remarkably, almost all FZNF genes are located on the long arm of chromosome four, a region previously noted for its enrichment in TEs, ZNF and immunity genes, as well as a wide variety of small RNA-encoding loci (Howe et al., 2016; Howe et al., 2013). Across the cyprinid family, the copy number of predicted FZNF genes varies substantially, even within closely related species (Fig. 1B); within the *Danio* genus alone, the number of predicted FZNF genes varies by an order of magnitude. To further validate these copy number estimates, we used raw reads from four *Danio* species and compared the read coverage over FZNF genes to coverage over BUSCO (Benchmarking Universal Single-Copy Orthologs) genes. In three out of four cases, there was good agreement between the two, but in the case of *Danio albolineatus*, the median read depth over FZNF genes was approximately 50% greater than that of BUSCO genes (Supp. Fig. 3A). This indicates that in many species, the true number of ZNF genes is likely to be far higher than gene annotation pipelines can predict, highlighting the difficulty of assembling and accurately counting these highly repetitive genes.

Finally, to verify that our automated ZNF ORF counts are good proxies for the number of annotated ZNF genes in a species, we compared these counts to those of our reannotated FZNFs, as well as independent estimates of KZNF copy number in tetrapods (Imbeault et al., 2017). In both cases, we found good agreement between the simpler ORF counts and higher quality estimates based on gene annotations (respectively, Spearman’s rho = 0.91, p-value = 2.6e-59; rho = 0.97, p-value = 1.7e-05; Supp. Fig. 3B, C). This relationship is approximately linear, but ORF counts grow faster than annotated gene counts due to factors such as genes having multiple exons and the presence of pseudogenes or gene fragments. Satisfied that our gene annotations were of sufficient quality for further downstream analysis, we began to test for evidence of coevolution between ZNFs and TEs.

### Correlation between ZNFs and retroelements

Early evidence for a coevolutionary relationship between TEs and ZNFs was reported by Thomas and Schneider, who observed a strong correlation between ZNF and retroelement copy number in a sample of 26 vertebrate species (Thomas and Schneider, 2011). By leveraging the greatly increased number of currently available genome assemblies, we sought to reproduce and expand on these results. Using the same process as for ZNF ORFs, we searched for reverse transcriptase and RNaseH domains (Supp. Data 1-2). These domains are highly conserved, and characteristic of proteins encoded by long terminal repeat (LTR) and non-LTR retroelements, making their counts rough proxies for the number of autonomous retroelement insertions in each genome.

A challenge with this approach is to address the issue of autocorrelation produced by the nonindependence of phylogenetically related species (Felsenstein, 1985). Since there is currently no well-established phylogenetic tree that includes all sequenced metazoan genomes, we opted to select a single species per taxonomic family, leveraging the fact that both ZNF and TE turnover is typically very rapid, so that even closely related species often have largely independent sets of both (Liu et al., 2014; Imbeault et al., 2017). This filtering procedure reduced the number of analyzed species to 828.

For this set of 828 genomes, we examined the correlation between the estimated number of ZNF and retroelement ORFs. This analysis revealed a significant positive correlation across all metazoans, as well as significant positive correlations across major phyla with at least 15 representative species, with the exception of nematode worms (Fig. 2A). Within smaller taxonomic groups, this relationship holds in most cases tested (Supp. Data 1), with some notable exceptions. For example, birds have lost most KZNFs (Imbeault et al., 2017), such that the median number of ZNF ORFs in birds is a quarter that of other chordates (121 vs 489); correspondingly, there is no correlation between ZNF and retroelement copy number. Similarly, we observe no correlation across dipteran flies, which possess ZAD-ZNFs – these have been studied in some depth in *Drosophila melanogaster* and have diverse roles in heterochromatin organization, but do not appear to specifically target TEs for silencing (Kasinathan et al., 2020; Chung et al., 2007; Baumgartner et al., 2022). Notwithstanding these exceptions, our analysis indicates that ZNF and TE copy numbers are positively correlated across the breadth of metazoans.

To further control for phylogenetic inter-dependence, we focused on the FZNF family in cyprinid fish, making use of our refined gene annotations and improved estimates of genomic TE content generated with dnaPipeTE (Goubert et al., 2015). Using these more accurate annotations, we find that the correlation between ZNFs and TEs persists (Fig. 2B). Importantly, this relationship holds when explicitly controlling for species relatedness by comparing phylogenetically independent contrasts in FZNF copy number and TE content (Fig. 2C).

**Figure 2.**
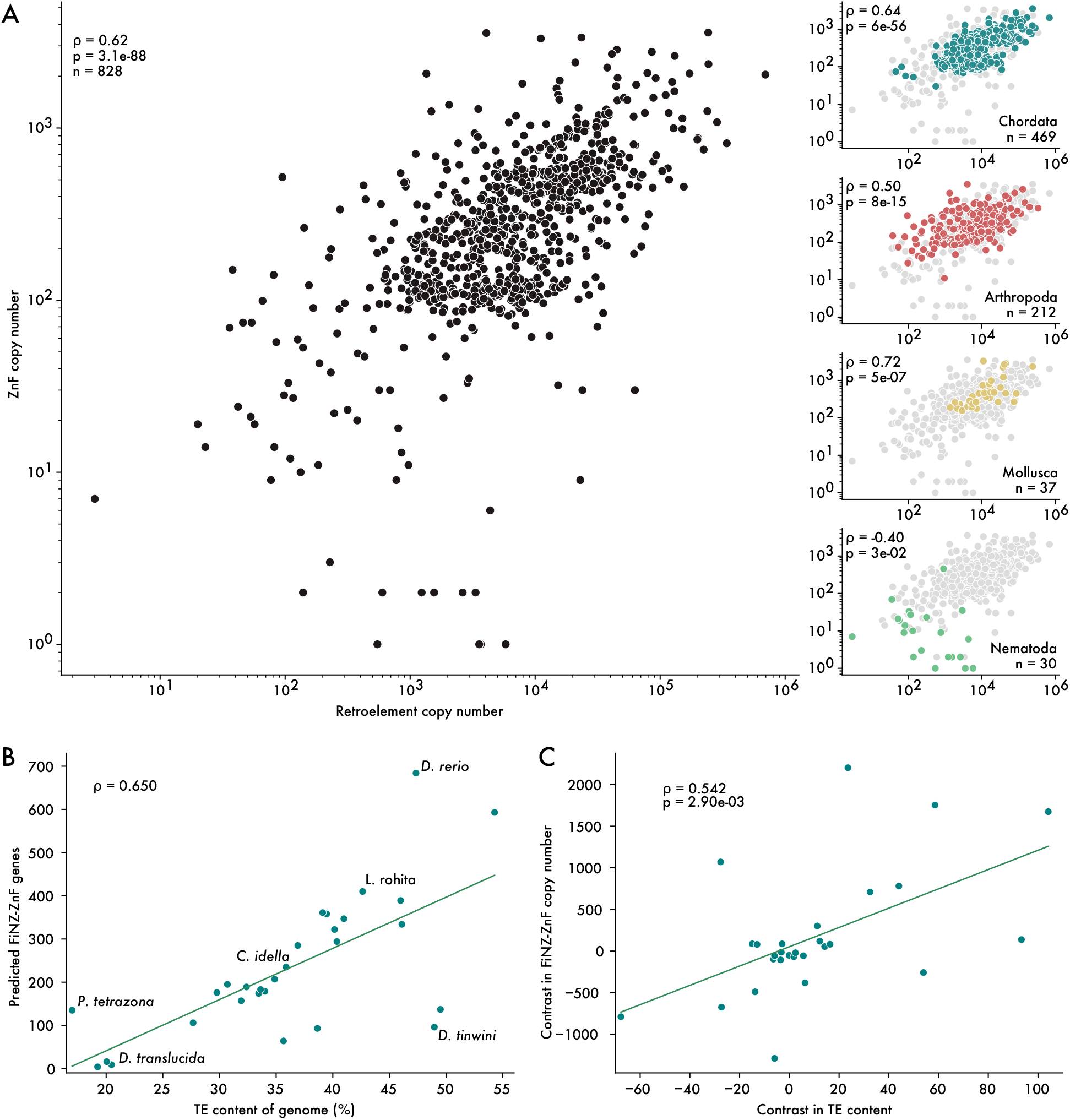
ZNF copy number correlates with genomic TE content. A) Spearman’s rank correlations between estimated number of retroelement insertions and ZNF ORFs. Each point is a representative species from a taxonomic family. Subplots on right hand side show correlations for individual phyla. B) Uncorrected Spearman’s rank correlation between TE content and FZNF copy number in cyprinid fish. C) Spearman’s rank correlation on phylogenetically independent contrasts.

Studies of the relationship between KZNFs and TEs thus far have been retroelement-centric, since widely used mammalian model organisms typically have relatively few DNA transposons (Platt et al., 2018). In contrast, cyprinid fish have a wide diversity of different TE classes, and particularly DNA transposons (Chang et al., 2022; Chen et al., 2019; Howe et al., 2013; Xu et al., 2014). We therefore asked whether the correlation between cyprinid FZNFs and TEs held across different TE classes. It is not clear whether a correlation with DNA transposons is expected, since these are typically less transcriptionally active than retroelements, at least in zebrafish (Chang et al., 2022), and might not be targeted by ZNFs. Yet, we found that the correlation was actually the strongest for DNA transposons (Supp. Fig. 4; rho = 0.6, p = 4e-04, phylogenetically independent contrasts), and in the case of LINEs there was no significant correlation. These results suggest that the relationship between FZNFs and TEs is not solely driven by transcriptionally active retroelements but applies to a broad range of TEs.

There are two non-mutually exclusive explanations for the correlation between TEs and ZNFs. First, ZNFs may physically interact with TEs, as is the case with KZNFs (Imbeault et al., 2017; Ecco et al., 2017). Second, the TE content of the genome may have an enhancing effect on the rate of gene duplication, such that genomes with large TE content will tend to have larger gene families. The latter model is conceivable because TE accumulation is thought to promote ectopic recombination – and therefore gene duplication or deletion – in a length- and copy number-dependent fashion (Charlesworth and Langley, 1989; Tollis and Boissinot, 2012; Jurka et al., 2004). Under this latter model, one would predict that TE content would positively correlate not only with the copy number of ZNFs but also that of other expansive gene families. To explore this possibility, we used the same approach as used to count ZNFs to estimate the copy number of mammalian olfactory receptors. Olfactory receptors are membrane proteins known to turn over rapidly across species but are not expected to interact physically with TEs. In a sample of representative species from 103 mammalian families, we observed a non-significant correlation between the number of retroelement and olfactory receptor ORFs (Spearman’s rho = 0.12, Supp. Fig. 5). While this analysis does not rule out the possibility that TE content has an impact on the rate of gene duplication, it does imply that the effect is insufficient to explain the correlation observed between ZNFs and TEs.

### ZNFs are predicted to bind TE sequences

Based on our findings thus far, we reasoned that ZNF proteins are most likely interacting with TEs through sequence-specific recognition of their DNA sequences (although occasionally ZNFs also bind RNA (Font and Mackay, 2010)). We therefore sought to test whether ZNF binding motifs are enriched in TE sequences. Defining the binding specificity of individual ZNF genes experimentally is labor-intensive, but computational methods have been developed to predict ZNF binding motifs directly from their protein sequence (Persikov and Singh, 2014; Najafabadi et al., 2015; Molparia et al., 2010; Kaplan et al., 2005). The most recent of these methods predicts binding motifs for individual ZNF domains using a random forest classifier (Najafabadi et al., 2015). We used this approach to predict DNA binding motifs for all ZNF ORFs from seven metazoan species with curated libraries of TE consensus sequences: four species with a relatively large repertoire of poorly characterized ZNFs (octopus, zebrafish, rice weevil and sea urchin), two “positive control” species known to use ZNFs to target TEs (human and mouse), and an expected “negative control” species, *Drosophila melanogaster*, whose ZNFs are apparently not targeting TEs directly (Supp. Data 3).

First, we used previously published ChIP-exo data for human KZNFs (Imbeault et al., 2017) to confirm that computationally predicted DNA binding motifs were similar to those experimentally determined. Searching for each of 236 experimentally determined KZNF binding sites in the set of predicted binding motifs, we found that 11 (5%) produced significant matches to their predicted counterpart (q-value < 0.05), compared to 0 matches after shuffling the predicted motifs. Similar results have been demonstrated by others working with mice (Wolf et al., 2020). As a second positive control, we compared the predicted binding motifs for human and mouse ZNF ORFs to libraries of their consensus TE sequences and observed a significant enrichment of predicted matches within TEs, compared to matches using shuffled motifs (Fig. 3A). This result is expected given that KZNFs are known to target TEs and confirms that predicted binding sites recapitulate, at least in part, experimentally obtained motifs. For example, we were able to recapitulate human ZNF320 binding to LTR14a and ZNF483 binding to L1PA7 (Supp. Fig. 6). These controls demonstrate that while computational prediction of ZNF binding specificity is far from solved, it is sufficient to capture a proportion of biologically relevant binding activity.

**Figure 3.**
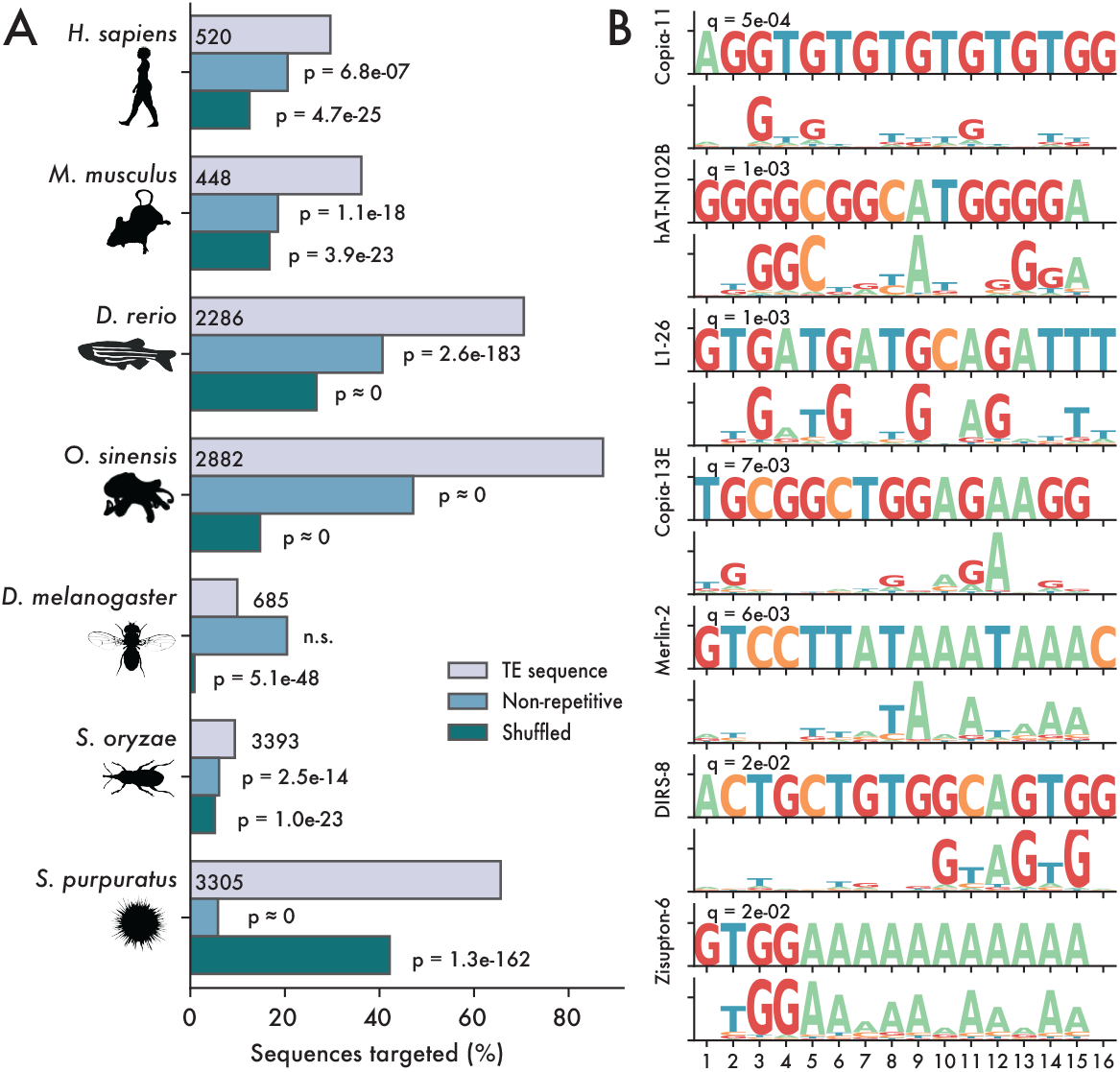
ZNFs are predicted to bind TE sequences. A) Percentage of sequences matched by predicted ZNF motifs, with a q-value cutoff of 0.05. Grey bars represent matches to libraries of consensus TE sequences, while blue bars represent matches in equally sized libraries of non-repetitive genomic sequence. Green bars represent shuffled motifs, searched against consensus TE libraries. P-values are calculated as one-tailed binomial tests for the probability of observing at least x matches in the TE library, using binding frequency to non-repetitive sequences or with shuffled motifs as a base probability. B) Selected matches between Danio rerio consensus TE sequences and ZNF ORFs.

We therefore turned our attention to the four species whose ZNFs are less studied (zebrafish, octopus, rice weevil, sea urchin), predicting binding motifs for all ZNF ORFs within each species and examining the frequency of these motifs in consensus TE sequences derived from each respective species (Fig 3A). Using a false discovery threshold of q = 0.05, we searched for significant matches between our predicted ZNF binding motifs and TE consensus sequences, then compared this to equivalently sized libraries of non-repetitive genomic sequence. For all four species, we observed a significant enrichment of matches to the TE sequences compared to non-repetitive sequences (Fig 3A). By contrast, we do not observe such enrichment for *Drosophila melanogaster*. Rather, for that species we found a greater percentage of non-repetitive sequences matched by predicted ZNF binding motifs than expected from shuffled motifs (Fig 3A). This finding is consistent with previous functional studies of *Drosophila* ZAD-ZNFs (Kasinathan et al., 2020) and our observation that the copy number of dipteran ZNFs does not correlate with the number of TE copies (Supp. Data 1). Notably, we also found that in all species both retroelements and DNA transposons were predicted to be targeted by ZNFs (Supp. Data 3).

As an additional control, we repeated the analysis using shuffled motifs, and found in all species that the predicted motifs bound a higher percentage of TE sequences compared to the shuffled motifs (Fig. 3A). We note, however, that shuffled motifs still recognize a substantial fraction of sequences in most species, likely due to the fact that ZNFs are often predicted to bind simple sequence repeats or homopolymeric tracts that would be unaffected by shuffling (e.g. Fig. 3B). Finally, to test species specificity, we searched the *Danio rerio* TE library using human ZNFs and compared these to zebrafish ZNFs. We found that zebrafish ZNFs were far more likely to target zebrafish TEs than were human ZNFs (odds ratio = 29.5, p ≈ 0.0, Fisher’s exact test). These results suggest that the preferred binding of ZNFs to TE sequences is not unique to mammalian KZNFs but likely a general property of many metaozan ZNF families.

### FZNFs are under positive selection in zebrafish

TEs evolve rapidly and are under strong selection to evade repression by their hosts; many KZNF proteins are in direct competition with TE families, leading to signatures of arms races between the two (Bruno et al., 2019; Fernandes et al., 2018; Jacobs et al., 2014). One such signature is positive selection (i.e. adaptive evolution) acting on the DNA contacting residues of ZNFs, i.e. those involved in TE recognition (Emerson and Thomas, 2009). We therefore tested for evidence of positive selection in *Danio rerio* FZNFs by comparing rates of synonymous and non-synonymous substitutions (dN/dS). Values of dN/dS > 1 denote an excess of non-synonymous changes and suggest that natural selection accepts novel amino-acid changes at a rate faster than expected under a neutral model.

In practice, it is challenging to confidently align ZNFs due to their repetitive nature and tendency to undergo internal rearrangements and gene conversion, and errors in this process can lead to false positives in assessment of dN/dS ratios (Mallick et al., 2009). To account for this, we restricted our analyses to seven clades of recently duplicated paralogs unique to *Danio rerio* (with a minimum of ten members - see methods for details). Using PAML (Yang, 2007), we performed likelihood ratio tests to compare two models of evolution for each clade – one in which the genes were evolving under purifying selection, and one in which some sites were assumed to be under positive selection (M2 vs M2a). In four of these seven clades, the model featuring positive selection (M2a) was significantly favored (p-value < 0.01, likelihood ratio tests, Supp. Data 4). Furthermore, in two clades we found that base-contacting residues of ZNF domains were significantly enriched for values of dN/dS > 1 (odds ratio = 2.45, p-value = 0.01; odds ratio = 2.84, p-value = 0.02; Fisher’s exact test).

As an alternative to dN/dS analyses, which are dependent on accurate gene alignments, we calculated the sequence entropy at positions in the canonical ZNF domain for each species in Fig. 3. The results revealed a clear trend for base-contacting residues to be amongst the most variable in sequence, with the possible exception of *Drosophila melanogaster* (Supp. Fig. 7). These data, combined with the enrichment of ZNF predicted binding sites in TE sequences, suggest that zebrafish ZNFs, and likely those of many other species, are evolving adaptively to target specific TE families.

### FZNFs are expressed in distinct waves during embryogenesis

TEs – retroelements especially – are highly active during metazoan embryogenesis, as this provides them with the opportunity to transpose in the cells giving rise to the adult germline, thus ensuring the vertical inheritance of new insertions (Haig, 2016; Ansaloni et al., 2019; Rodriguez-Terrones and Torres-Padilla, 2018; Chang et al., 2022). Previous work has shown that zebrafish ZNFs are expressed at the onset of zygotic genome activation (ZGA), mirroring the pattern of KZNFs in human and mice, and consistent with a potential role in TE silencing (Pontis et al., 2019; White et al., 2017; Hadzhiev et al., 2019). We sought to further explore the timing of FZNF expression, predicting that if they are deployed to repress TEs, their expression should overlap with that of TEs. Using our de-novo FZNF annotations, we remapped previously published RNA-sequencing data covering the first 24 hours of zebrafish development (White et al., 2017). Setting a lower limit of 0.5 transcripts per million (TPM) to call genes as expressed, we find that approximately half of all FZNFs (306 out of 684) are expressed during early development.

Recently published work identified two distinct waves of ZNF expression in zebrafish, named “sharp peak” and “broad peak” (Hadzhiev et al., 2019; Hadzhiev et al., 2021). Sharp peak FZNFs share a distinct promoter architecture consisting of a clearly identifiable TATA-box and are expressed in the minor wave of ZGA, followed shortly after by TATA-less broad peak ZNFs in the major wave. With our updated FZNF annotations, we recapitulated these findings, observing expression of 80 sharp peak and 204 broad peak genes, hereafter termed “dome”, and “late” based on the timing of their expression (Fig. 4A). The timing of these peaks overlaps with that of most TE families, and in the case of dome stage FZNFs, precedes almost all zygotic TE transcription.

**Figure 4.**
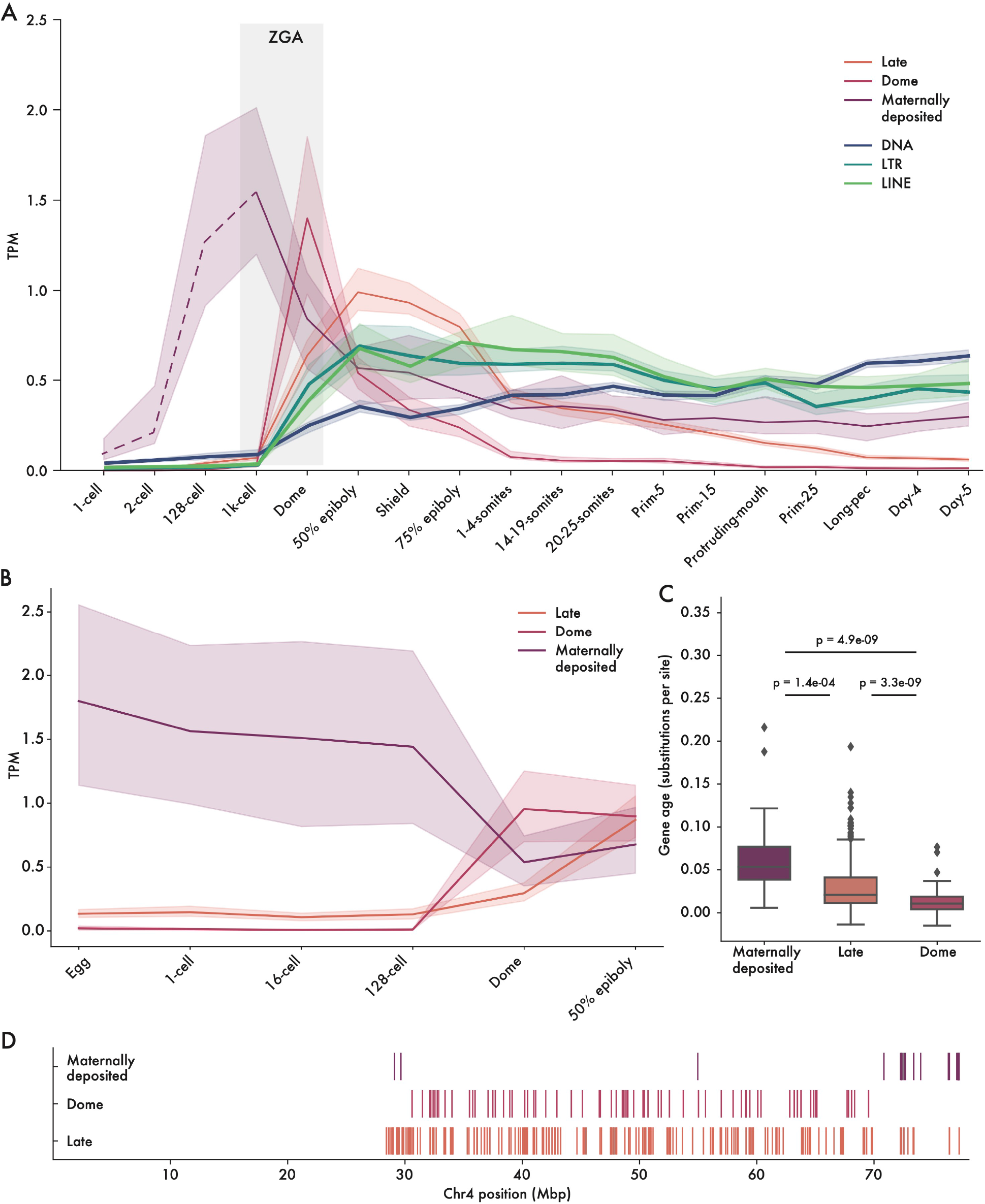
Embryonic expression of FZNF genes in zebrafish. A) Median expression trajectory of 306 genes with expression of at least 0.5 TPM in at least 1 stage, separated by cluster. Shaded borders represent 90% confidence intervals. Light grey bar shows approximate timing of ZGA. Expression trajectory of maternally deposited FZNFs prior to 1000-cell stage is dashed to reflect the fact that poly-A selected RNA sequencing is not an accurate reflection of true mRNA levels prior to ZGA. B) FZNF expression from rRNA-depleted reads, prior to ZGA. C) Maternally deposited genes are significantly older, whilst those expressed specifically at Dome stage are young, and generally unique to D. rerio. Ages were quantified as branch lengths on a neighbor-joining phylogenetic tree generated from pairwise distances. D) The majority of maternally deposited FiNZ genes are located in a specific, sub-telomeric cluster. Each vertical line represents the midpoint of an expressed FZNF gene.

We also found a small subset of approximately 22 FZNF genes with maternally deposited transcripts (Fig. 4A). To our knowledge, these have not previously been described, likely due to their incomplete annotation in Ensembl and RefSeq gene sets. Furthermore, maternally-deposited mRNAs in zebrafish and other animals are, initially, only partially poly-adenylated (Winata et al., 2018; Cui et al., 2013; Potireddy et al., 2006), and are therefore difficult to detect when using polyA-selected RNA-seq platforms. As the White et al. data falls into this category, we remapped rRNA-depleted reads from a second dataset covering early embryogenesis (Winata et al., 2018), which confirmed that these maternally-deposited transcripts are indeed present in both the egg and 1-cell zygote (Fig 4B).

Given the markedly different expression pattern of maternally deposited FZNFs, we looked for other features differentiating them from their zygotically expressed counterparts. While the majority of *Danio rerio* FZNFs are recently duplicated, particularly dome stage genes, those that are maternally deposited are conserved across species and are significantly older (Fig. 4C, Supp, Data. 5). Moreover, they are physically co-localized in the subtelomeric region of Chr 4q (Fig. 4D), which lacks the repeat density that characterizes much of Chr 4q (Chang et al., 2022). The age of maternally deposited genes, as well as their degree of species conservation, is hard to reconcile with a role in TE targeting, as TEs turn over rapidly and are frequently species-specific. Rather, the deeper conservation and physical clustering of these maternally deposited FZNFs is suggestive of a role in development that differs from that of Dome and Late-stage FZNFs.

### FZNFs repress LTR retroelement expression during early development

Next, we turned to the function of FZNF genes. Based on their embryonic expression, predicted binding specificity and strong correlation with TE content, we predicted that the bulk of FNZF genes play a role in TE silencing. If true, then knocking down their expression ought to lead to a corresponding increase in TE expression. To test this, we employed a Morpholino oligonucleotide strategy aimed at blocking translation of most FZNF proteins simultaneously. By aligning the first exon of the FZNF annotations from zebrafish, we were able to design a Morpholino predicted to target the translation start site of 447 out of 684 annotated FZNFs (Fig. 5A). We injected zebrafish embryos either with this Morpholino or a scrambled control, then compared gene and TE expression using RNA-seq. Embryos were collected for sequencing at shield stage, as this corresponds with a period at which many TE families and most FZNFs are robustly expressed (Fig. 4A).

**Figure 5.**
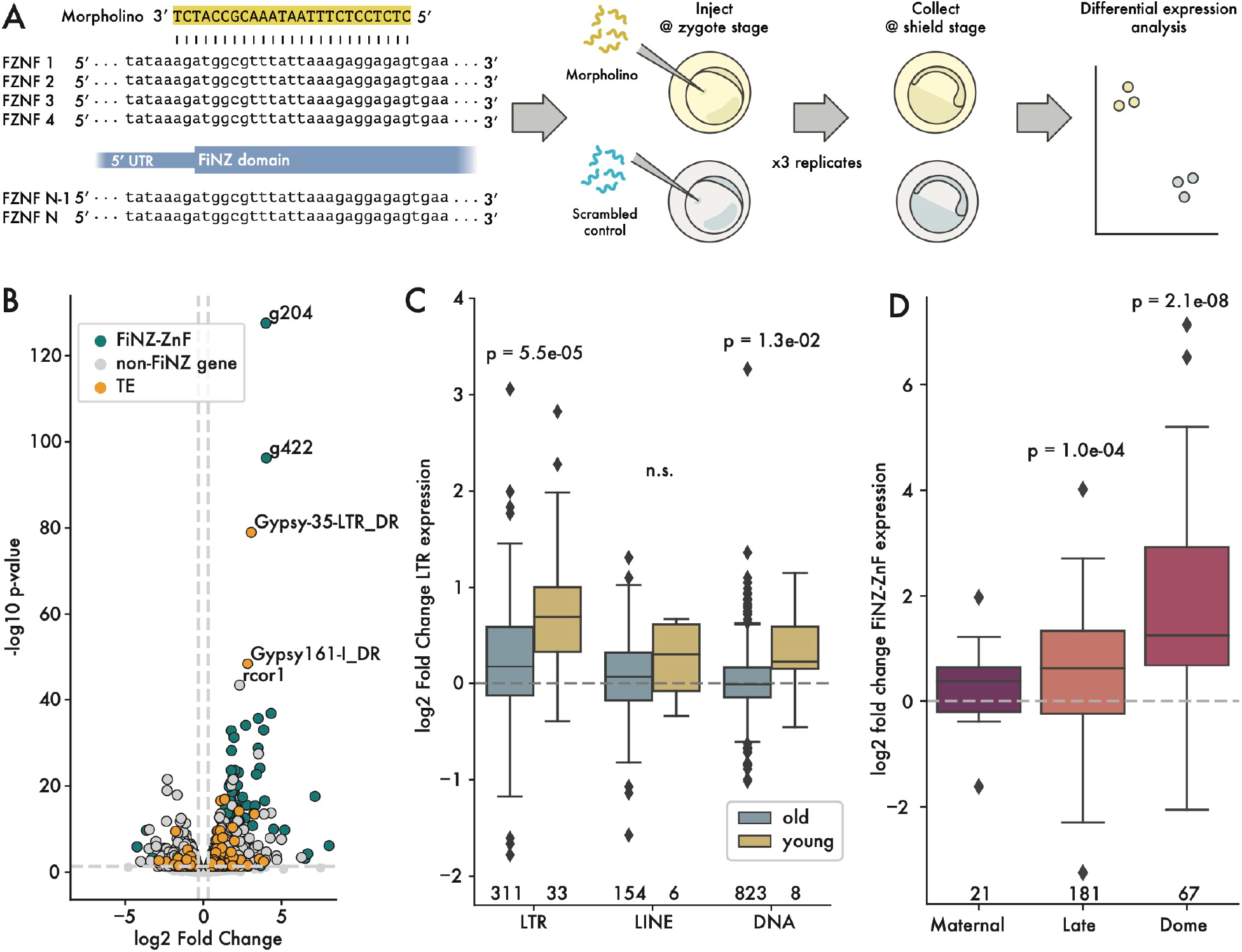
Up-regulation of LTR retroelements in response to FZNF knock down. A) Cartoon showing design of Morpholino-based, translation-blocking FZNF knock-down experiment. B) Log2 fold change vs p. value, comparing FiNZ translation blocking morpholinos to scrambled morpholino control. C) Log2 fold change comparison between old and young TE families, with “young” being defined as those whose median insertion age was less than 0.01 substitutions per site. P-values are Wilcoxon rank-sum tests comparing old and young in each TE class. D) Log2 fold change of different FZNF categories. P-values are binomial tests for each category, assessing likelihood of seeing at least x upregulated genes, assuming a null hypothesis where there is a 50% chance of being either up- or down-regulated. Numbers below boxes in C) and D) give number of data points per box.

After sequencing and read mapping, we carried out quality controls before comparing TE expression between treatment and control groups. First, we performed principal component analyses of batch-corrected samples, which showed clear separation between the FZNF knock-down and control groups (Supp. Fig. 8A). Using a false discovery rate threshold of 0.05 and an absolute log2 fold change of at least 0.32 to call differentially expressed genes and TE families, we observed a total of 999 differentially expressed genes, excluding FZNFs and TEs (Supp. Data 6). Amongst these, a GO-term enrichment analysis revealed no notable enrichment for biological processes beyond general terms related to embryonic development, metabolic processes and translation (Supp. Data 7). Importantly, we did not observe any enrichment for stress response pathways, nor did we observe a significant change in *tp53* levels (log2 fold change = −0.01), which has been known to cause off-target effects in Morpholino knockdowns (Bedell et al., 2011). Overall, there was no significant difference in the number of up- vs down-regulated non-FZNF genes (p = 0.02, binomial test).

As the expression of TEs and FZNF genes is highly dynamic during the first 24 hours of zebrafish development (Chang et al., 2022; Laue et al., 2019; Guo et al., 2021), we sought to ensure that the embryos were staged appropriately, as differences in the relative stage of FZNF knockdown and control groups could lead to artefactual observations of differential expression. We therefore compared our samples with samples from White et al., covering 50% epiboly, shield stage, and 75% epiboly (White et al., 2017). Treating these samples as additional control groups, and after batch correction, we found that our samples clearly clustered with shield stage samples from White et al. (Supp. Fig. 8B, C).

We then asked whether TE expression was up-regulated in the FZNF knock-down group. Looking at the overall number of differentially expressed TE families, we found that TEs were significantly more likely to be up-regulated than down-regulated, consistent with de-repression following FZNF knockdown (odds ratio = 3.78, p-value = 6.1e-08, Fisher’s exact test; Fig. 5B, Supp. Fig. 8D-E). Separating by TE class, we found that this trend was driven predominantly by LTR elements (odds ratio = 6.56, p-value = 1.2e-08, Fisher’s exact test; Supp. Fig. 9A). Since LTRs include many of the youngest TE families in the zebrafish genome (Chang et al., 2022), we reasoned that their enrichment amongst up-regulated genes and TE families may reflect a tendency for young, active families to be most intensively targeted by FZNFs. We therefore compared young and old TE families to each other, defining the former as those whose insertions averaged fewer than 0.01 substitutions per site (Chang et al., 2022). This analysis revealed that the magnitude of de-repression following FiNZ-knockdown was significantly greater amongst young families (p = 5.5e-05, Wilcoxon rank sum test), and that this trend held across different TE classes (Fig. 5C). Thus, FZNF knockdown leads to significant de-repression of young, potentially active TE families.

In addition to age, we also looked at the timing of expression, since by collecting at “shield” stage (approximately 6 hours post fertilization), we would not be able to observe the effects on those TEs whose expression peaked at later time points. Using hierarchical clustering to separate TEs into groups with expression peaks either before or after shield stage, we found that those TE families which were expressed earlier were significantly more up-regulated than those expressed later (Supp. Fig. 9B).

Curiously, the genes most up-regulated by FZNF knockdown were FZNFs themselves, for example, g204 and g422 (Fig. 5D). While we cannot currently rule out the possibility that the Morpholinos inhibit the degradation of their target sequences, to our knowledge, such effects have not previously been reported in the literature. Alternatively, these results suggest that FZNF genes are themselves targeted for repression by FZNF proteins, thus forming a negative feedback loop. In this way, their expression would be robustly switched off once sufficient FZNF protein had been generated, without the need for additional regulatory control. Such negative feedback loops have recently been documented for mammalian KZNFs (Pontis et al., 2022).

## Discussion

In this work, we observed a deep coevolutionary relationship between TEs and ZNFs that spans the breadth of metazoans. This is most clearly seen in the strong positive correlation between TE and ZNF copy numbers, which persists across at least 750 million years of animal evolution and is independent of the various accessory domains that characterize different ZNF families. We also found that ZNFs from a range of metazoans are predicted to bind TE sequences, and in the case of FZNFs in zebrafish, are evolving under positive selection in ZNF arrays. Finally, through simultaneous knock-down of a large subset of FZNFs during zebrafish embryogenesis, we demonstrated that this family represses young, recently active TEs. Since the FiNZ domain is unique to cyprinid fish, which lack both the KRAB domain and its cofactor KAP1 (a.k.a. TRIM28), the silencing abilities of FZNFs and KZNFs appear to have evolved convergently. This suggests that the role of ZNFs in TE targeting and silencing is driven by the intrinsic plasticity of ZNF DNA binding activity, as opposed to being a function of specific accessory domains.

The extensively duplicated ZNF families of metazoan species are rarely observed in other eukaryotic taxa, where instead, small-RNAs acting in concert with Argonaute proteins are the primary drivers of TE heterochromatinization (Hall et al., 2002; Volpe et al., 2002; Allshire and Madhani, 2018; Heger et al., 2020; Najafabadi et al., 2017). Thus, ZNF-based TE targeting is likely to be a metazoan innovation. However, there are several metazoan groups whose ZNF families appear to behave atypically. For example, we found no correlation between ZNFs and retroelements in dipteran flies, which includes intensively studied species such as *Drosophila melanogaster* and *Anopheles gambiae*, nor did we observe a binding preference for TEs in *Drosophila* ZNFs. However, in *Drosophila* the piRNA system (itself a branch of the Argonaute-siRNA system) is directly involved in pre-transcriptional silencing of TEs during embryogenesis, with *piwi* knockdowns resulting in a failure of H3K9me3 to establish over otherwise silenced TE loci (Sienski et al., 2012; Wei et al., 2021; Fabry et al., 2021). Less is known about mosquitoes, but it is clear that they have undergone numerous duplications of *piwi* homologues, and have a complex suite of piRNAs that includes many arising from non-TE satellite sequences (Ma et al., 2021; Gamez et al., 2020). Similarly, nematode worms, which typically have few ZNF genes, have undergone extensive diversification of Argonaute genes (Buck and Blaxter, 2013). While research into the function of these Argonaute homologs remains in the early stages (Seroussi et al., 2022), both the dipteran and nematode cases point to potential functional overlap between the Argonaute/piRNA system and ZNFs.

It is still an open question whether silencing of TE transcription *per se*, is the primary function of metazoan ZNFs as a form of genome defense system. While depletion of both KZNFs and FZNFs (this study) have now been shown to unleash TE expression, there are many paradoxical observations that challenge the idea of ZNF as an adaptive defense system against TE invasion and suggest transcriptional repression itself may be secondary to other functions. First and foremost, it appears that ZNFs evolve at a glacial pace relative to the speed at which novel TE families can establish and propagate within a species. For example, the DNA transposon P-element spread throughout all wild populations of *Drosophila melanogaster* in the space of approximately 30 years, and the retrovirus KoRV has been spreading in koalas exogenously and endogenously over a similar period (Anxolabéhère et al., 1985; Tarlinton et al., 2006); in contrast, observations in primates imply that the emergence of novel KZNF repressors takes place over millions of years (Imbeault et al., 2017; Ecco et al., 2017). On the other hand, the piRNA 1 system is able to adapt much more rapidly, as TE insertions themselves form the raw material for novel piRNAs – indeed, in the case of both KoRV and P-element, piRNAs have already emerged to mitigate their deleterious effects (Brennecke et al., 2008; Yu et al., 2019). Thus, with respect to their ability to quickly respond to novel TEs, the piRNA system appears better suited as a front-line defense than ZNFs.

In addition to the speed at which they evolve, other features of ZNFs are also difficult to reconcile with a general function in repressing TE transcription. While the majority of KZNFs are expressed during early embryogenesis and specifically target TEs, approximately a third target non-TE sequences, such as satellite repeats, gene promoters, and even other KZNFs (Imbeault et al., 2017). There are also examples in both mouse and human of conserved clusters of KZNFs that target TEs younger than themselves, implying they had targets predating the emergence of their current TE families (Imbeault et al., 2017; Wolf et al., 2020; Wolf et al., 2015). These observations have led others to propose that, rather than simply repressing TEs, KZNFs play an important role in their domestication by taming their cis-regulatory activities and facilitating their integration into existing gene regulatory networks(Ecco et al., 2017; Pontis et al., 2019; Bruno et al., 2019). Under this model, KZNFs would act as tolerogenic agents promoting the evolution of species-specific gene regulatory networks. While this is an attractive model, it is important to clarify that this feature of KZNFs cannot be the initial driver of their rapid evolution and turnover, since TE domestication is a process that takes place over many generations, whereas selection primarily acts on currently extant individuals (Sniegowski and Murphy, 2006).

We therefore propose a non-mutually exclusive alternative to the transcriptional repressor hypothesis, in which metazoan ZNFs instead serve as genome stabilizers, protecting DNA from ectopic recombination by nucleating heterochromatin to repetitive regions of the genome. Heterochromatin has a strong suppressive effect on homologous recombination and prevents the formation of gross chromosomal rearrangements (Allshire and Madhani, 2018; Grewal, 2010). For example, histone H3K9 methylation prevents repeat- and/or TE-mediated recombination and replication stress in worms and fission yeast (Zeller et al., 2016; Sasaki et al., 2010). Importantly, the propensity of TEs to cause deleterious chromosomal rearrangements via non allelic recombination is thought to be the primary factor preventing their accumulation in genomes (Allshire and Madhani, 2018; Sasaki et al., 2010; Kent et al., 2017; Tollis and Boissinot, 2012). Thus, the deployment of sequence-specific ZNFs to deposit heterochromatin over discrete TE families might explain in part why some organisms can sustain large-scale amplification of TE families as seen in a subset of animal genomes such as octopus, weevils, sea urchin, cyprinid fish, and mammals to name a few highlighted in this study. If genome stabilization is the principal function of most metazoan ZNFs, then it relieves the requirement for rapid adaptation to novel TEs, since the effect of TEs on rates of non-allelic homologous recombination are not felt until the TEs in question have reached significant copy number. Similarly, it would explain why satellite sequences or non-autonomous DNA elements, which do not need to be transcribed to amplify, appear to be targets of many ZNFs.

Much remains to be discovered about the role of ZNF genes in metazoan evolution, but with the wealth of high-quality genome assemblies and improved tools for manipulating gene expression in a variety of organisms, it is now possible to answer many of the long-standing questions about this extraordinary gene family. In this work, we have demonstrated that the co-evolutionary relationship between TEs and ZNFs extends well beyond KZNFs and is likely a metazoan innovation for TE silencing and/or stabilization. In zebrafish, FZNFs have independently expanded to repress young, recently active TE families during early embryonic development. Across animals, there are dozens of other such uncharacterized ZNF families, many of which may reveal exciting new biology, and which have potential uses in the development of genome engineering tools.

## Methods

### Predicting ZNF ORF copy number

Initial estimates of ZNF ORF copy number were achieved by extracting all possible ORFs from 3,221 representative metazoan genomes available from the NCBI assembly database, as of 2021-12-06. To facilitate this process, we set minimum and maximum ORF sizes of 375 and 10,000 base pairs, respectively. The resulting amino-acid sequences were searched for potential C2H2 ZNF domains using hmmsearch command line tool from the HMMER suite (Eddy, 2011), and the PFAM “zf_C2H2” (PFAM: PF00096) HMM as a query profile (Mistry et al., 2021). Resulting hits were merged if they were within 100 residues of each other, and filtered to exclude sequences with fewer than five ZNF domains in total.

### Annotation of FZNF genes

For improved annotations of FZNF genes in 32 cypriniform fish and a milkfish outgroup, we first identified genomic regions containing candidate FZNF genes using BLAST to search for matches with a consensus FiNZ domain sequence. This consensus sequence was generated from a set of *D. rerio* FZNF genes previously identified as being expressed during development (White et al., 2017). Using the “PPX” module of Augustus v3.3 (Hoff and Stanke, 2018; Keller et al., 2011), we carried out a first pass of the genomic regions identified in our set of cypriniform species, using a protein profile generated from the previously mentioned set of *D. rerio* annotations. To avoid biasing our search towards zebrafish, we used the results of this round of annotation to generate a new FZNF profile generated by sampling genes from all species. We then repeated the above procedure, and performed a final filtering step with HMMER (Eddy, 2011) to identify annotated genes containing both the FiNZ domain and ZNF domain (PFAM: PF00096).

For *D. rerio* specifically, we produced a separate, high-quality set of gene predictions by retraining Augustus specifically for FZNFs using a manually curated set of Ensembl predictions. This allowed us to generate predictions for 5’ and 3’ untranslated regions, for use downstream in analyses of gene expression. We explored the effect of including transcripts generated by Trinity(Grabherr et al., 2011) as hints for Augustus, but found that these reduced the quality of resulting annotations, likely as a result of the difficulty of assembling accurate transcripts for such repetitive genes. Full parameter details are available online.

### Predicting genomic TE content

We used two approaches to estimate genomic TE content. In the first, we focused on retroelements, using an identical approach to that which we used to estimate ZNF ORF number. In place of the PFAM zf_C2H2 domain, we instead used a selection of reverse transcriptase and RNaseH profiles (PF00078, PF07727, PF13456, PF00075, PF17917, PF17919). Resulting hits were merged as done for ZNFs, but no restrictions were placed on the number of hits per ORF. For more careful estimates of TE content for use when focusing on FZNF genes, we estimated the TE-derived proportion of genomes using dnaPipeTE(Goubert et al., 2015), a tool that offers considerable speed improvements over RepeatMasker/Modeller with comparable accuracy. This program requires short reads and genome size as input: for the former, we simulated reads using ART(Huang et al., 2012) and for the latter, we used genome assembly size as a proxy for true genome size.

### Cypriniformes phylogeny

To generate a phylogenetic tree of the Cypriniformes order (required to account for phylogenetic nonindependence in comparative genomic analyses) we first used the Actinopterygii-specific BUSCO database to extract intact single-copy orthologues from our set of 33 species. We then selected those protein sequences found in at least 10 out those species, producing a final set of 3,581 proteins. These were aligned separately using mafft v7.475 with parameters –globalpair and –maxiterate 1000 (i.e. ginsi) (Katoh and Standley, 2013) and trimmed to remove large insertions or poorly aligned regions using trimAl v1.4.rev15 with the –automated1 parameter set (Capella-Gutiérrez et al., 2009). All resulting files were combined to produce a concatenated super-gene alignment, which was used as input when generating a time-calibrated phylogeny.

IQ-TREE v2.0.6 was used to generate a maximum likelihood tree from a partitioned analysis of the supergene alignment (Minh et al., 2020; Chernomor et al., 2016; Hoang et al., 2018). Ultrafast bootstraps and Shimodaira–Hasegawa approximate likelihood ratio tests were used to determine branch support values, with 5000 and 1000 replicates respectively. A time-calibrated tree was generated using the integrated LSD2 module (To et al., 2016), constraining the date of the split between Gonorynchiformes and Cypriniformes (i.e. between *Chanos chanos* and all other species) to 162 Mya, based on a recent comprehensive phylogeny of teleost fish (Hughes et al., 2018).

### Predicting ZNF binding sites

To predict ZNF binding sites, we made use of the previously published tool, ZiFRC, with ZNF ORFs identified from our initial genome searches as input (Najafabadi et al., 2017; Najafabadi et al., 2015). To test for enrichment of predicted ZNF binding sites in various sequence sets, we used fimo and tomtom from the MEME suite of motif enrichment tools (v5.3.3) (Bailey et al., 2015). First, tomtom was used to compare our predicted motifs to motifs obtained from human ChIP-seq datasets (Imbeault et al., 2017). Following this positive control, we used fimo to calculate motif enrichment in three separate cases for each species.

In the first case, we searched our library of predicted ZNF binding sites against libraries of TE consensus sequences. For human, mouse, fruit fly and zebrafish, these libraries were taken directly from RepBase26.10, in the case of sea urchin and rice weevil, they were provided by direct communication with the authors of previously published studies on the TE content of these species (Panyushev et al., 2021; Parisot et al., 2021), and in the case of octopus, we generated a TE consensus sequence library using RepeatModeller v2.0.1 (Flynn et al., 2019).

For the second case, we searched a library of randomly selected non-repetitive genomic sequences, selected using bedtools v2.30.0 (Quinlan and Hall, 2010); this was done in a manner ensuring that, for each species, the resulting library size and distribution of sequence lengths was identical to that of the TE sequence library. In the third case, we searched the TE consensus sequences again, this time using shuffled binding motifs. Finally, we performed a cross-species comparison between zebrafish and human using fimo.

### Testing for positive selection

To test for evidence of positive selection in FZNF genes, we first searched for recently duplicated inparalogs. As multiple sequence alignments of ZNF genes can be unreliable, we calculated pairwise Needleman-Wunsch alignments between all pairs of 6,732 annotated genes from 29 cypriniform species with at least one FZNF gene, using the “needle” command line tool from the EMBOSS suite, v6.6.0 (Rice et al., 2000). The resulting pairwise distance matrix was used as input to generate a neighbor-joining phylogenetic tree, using the EMBOSS “fneighbor” tool. This gene tree was used to identify clades of recently duplicated in-paralogs within the cypriniform species tree. We selected seven clades from *Danio rerio*, where each clade contained at least ten members, and all members were within 30 amino-acids of each other in length. Each one of these clades was realigned with Prank, using a codon substitution matrix (Löytynoja, 2014; Kosiol et al., 2007), and all gaps in the resulting alignment were removed using using trimAl v1.4.rev15 (Capella-Gutiérrez et al., 2009). Finally, guide trees were generated for each clade using IQ-TREE v2.0.6 with a codon substitution model (Minh et al., 2020).

To test for evidence of positive selection, we used the “codeml” tool from the PAML4 software suite (Yang, 2007). We compared two sets of nested site models: first, M1a (NearlyNeutral) and M2a (PositiveSelection), and second, M7 (beta) and M8 (beta & ω). The transition to transversion ratio, κ, was calculated from the guide tree, while the dN/dS rate ratio, ω (i.e. the parameter being tested) was estimated from the data. Likelihood ratio tests with two degrees of freedom were used to compare M1a to M2a, and M7 to M8, respectively.

### Reanalysis of zebrafish developmental transcriptome data

To assess FZNF gene and TE expression, we used first used STAR (v2.7.5a) (Dobin et al., 2013) in multimapping mode to re-align reads from White et al. (White et al., 2017) to the *Danio rerio* GRCz11 reference genome, setting the maximum intron/read pair gap size at 500,000. To count reads from genes and TEs simultaneously, we used TEtranscripts-2.2.1’s TEcount tool in multi-mapper mode, with a GTF file combining Ensembl gene annotations with our FZNF annotations, along with TE locations as provided in Chang et al. (Chang et al., 2022). We used GTFtools v0.8.0 (Li et al., 2022) to calculate the median length of each gene, and used these measurements to calculate TPM for each gene/TE, allowing comparison of expression across stages. Specifically, for each transcript we calculate the sum of the union of exons. For genes, we then take the median length of transcripts associated with each gene, whereas for TEs we use the sum of all insertions in the genome, thus partially controlling for insertion copy number.

### Morpholino knockdown of FZNF translation

We ordered a translation blocking Morpholino with the sequence “CTCTCCTCTTTAATAAACGCCATCT” from Gene Tools LLC, which was sufficient to target 447 of our predicted FZNF mRNA transcripts with no mismatches. Wild-type, Tübingen strain zebrafish were maintained at 28°C on a 14/10hr light/dark schedule. Individual breeding pairs of male and female fish were separated overnight to induce naturally synchronized spawning and fertilization the following morning. To knock down FZNF expression, we injected 4.5ng of the custom Morpholino into fertilized *Danio rerio* zygotes, and in parallel, we injected embryos with the same quantity of a randomized control, consisting of a mixture of up to 4^25^ possible sequences. Embryos were collected at shield stage, and stored at −20°C immediately after collection. Each pair of injections was repeated in triplicate, such that 23 treatment and 19 control embryos were in the first batch, 30 treatment and 30 control in the second, and 31 and 30, respectively, in the third. Total RNA extraction was performed using the Qiagen RNeasy kit.

### RNA sequencing

We outsourced cDNA library preparation and sequencing to Novogene Inc. Samples were shipped in dry ice, and after quality controls, strand-specific libraries were prepared using poly-A enrichment. These libraries were pooled and sequenced in 150bp paired-end mode using the NovaSeq 6000 platform. After receiving sequencing data, we carried out quality assessment using FastQC (v0.11.9) (Andrews, 2010), and then mapped and counted reads according to the protocol previously described for reanalysis of White et al. data. Following this, differential expression analyses were carried out using the DESeq2 R package v1.26.0 (Love et al., 2014).

## Code and data availability

All scripts used in the generation and analysis of data for this project can be found at GitHub. Code relating to the broader analyses of metazoan ZNFs can be found at: https://github.com/jonathan-wells/metazoan-znfs, while code relating to FZNFs specifically can be found at: https://github.com/jonathan-wells/finz-znf. Supplementary data files containing processed data are provided, and all raw data is available upon request. Sequencing reads generated through this work will be available at NCBI SRA.

## Acknowledgements

We thank Dr. Joseph Fetcho for zebrafish husbandry support. We thank members of the Feschotte laboratory for valuable feedback and discussion. This work was supported by grant R35-GM122550 from the National Institutes of Health to C.F. J.W. is supported by a Human Frontier Science Program longterm fellowship (LT000017/2019-L). N.-C.C. is supported by a Distinguished Scholar Award from the Cornell Center for Vertebrate Genomics. C.C. was supported by a stipend from the Cornell Research Experience for Undergraduates program.

## Author contributions

J.W. performed all computational analyses. N.-C.C. performed FZNF Morpholino injections. J.M. carried out early studies of the FiNZ domain and provided general assistance with “wet-lab” experiments. C.C. generated and analyzed preliminary data from surveys of metazoan ZNF and TEs. N.R. carried out early annotation of *Danio* genus TEs and performed pilot wet-lab experiments. B.J. generated and analyzed preliminary data for testing correlation between TEs and other gene families. J.W. designed all experiments with input from N.-C.C., J.M. and C.F. J.W. wrote the manuscript and created figures with input from C.F. Initial project conception by J.W., J.M. and C.F.

