## Supplemental Figures for "Transposable elements drive the evolution of metazoan zinc finger genes"

**Supplementary Figures**

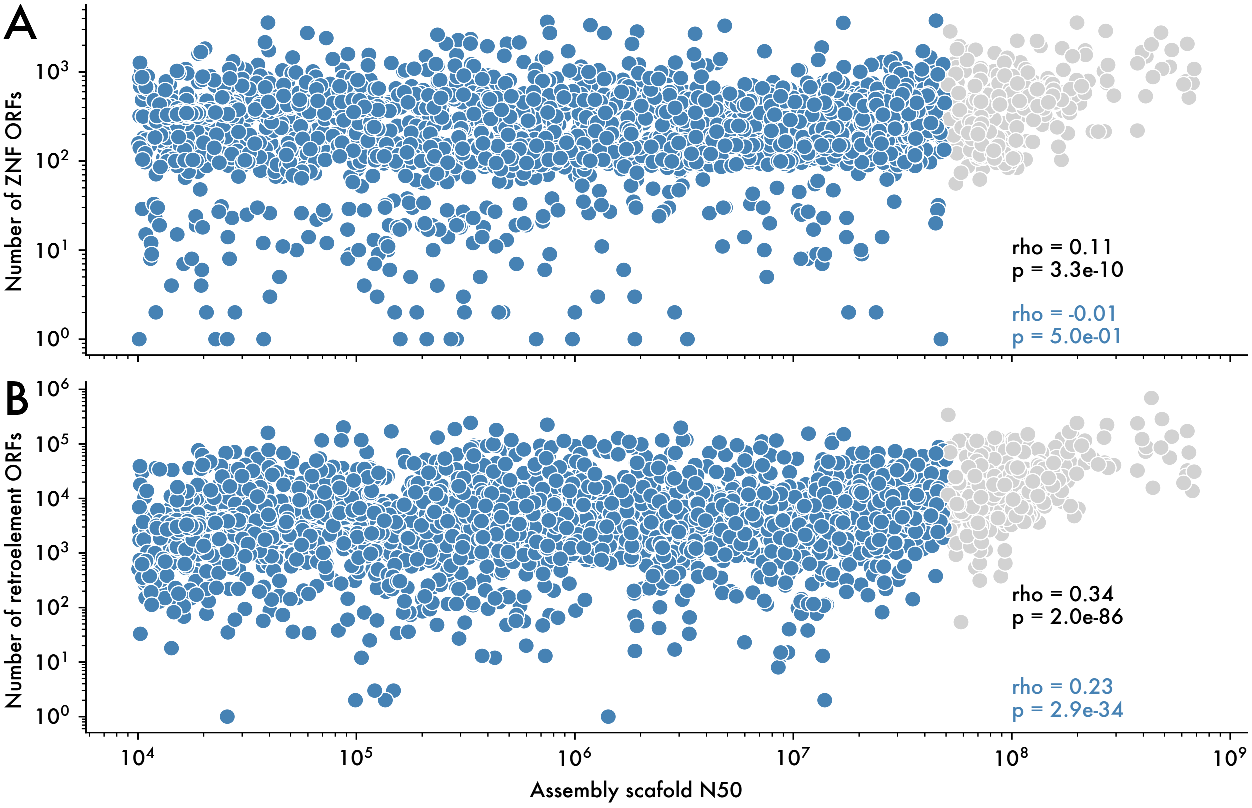


***Supplementary Figure 1. Correlation between scaffold N50 and retroelement and ZNF counts.*** *To assess the degree to which genome assembly quality biases our counts of ORFs, we compared scaffold N50 with A) ZNF and B) retroelement ORF copy number. This revealed low correlations, primarily driven by the fact that very large, high-quality genomes (and thus high scaffold N50 scores) almost always have high numbers of both retroelements and ZNFs.*

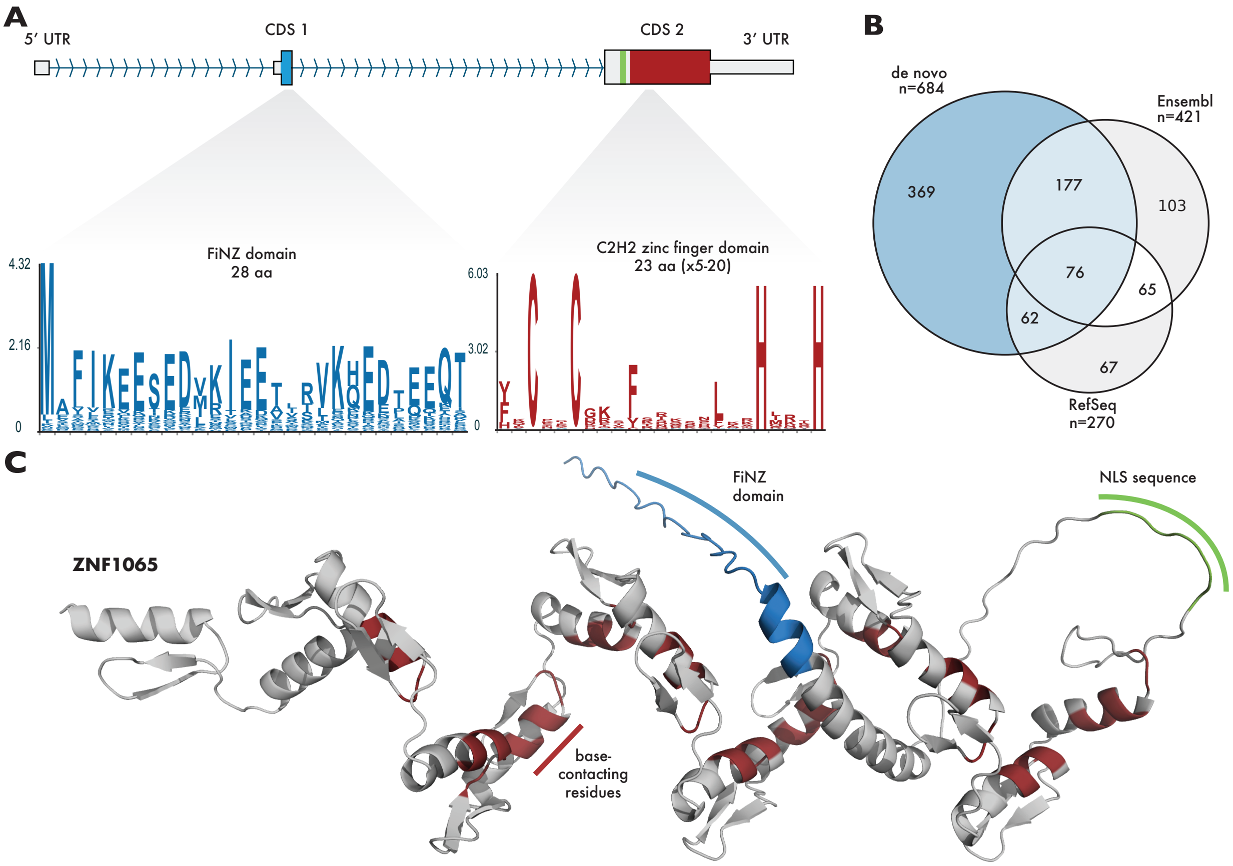


***Supplementary Figure 2. Typical structure and number of annotated FZNF genes in Danio rerio***
*A) The vast majority of FZNFs that we re-annotated in Danio rerio have two coding exons, the first containing the FiNZ domain itself, and the second containing a putative NLS sequence (green) and an array of tandemly repeated ZNF domains. B) Venn diagram showing overlap of different FZNF gene annotations. Annotations were categorized as matching if they overlapped over at least 70% of the gene body, ignoring untranslated regions. C) AlphaFold prediction of ZNF1065 (H0WEE1)* (Varadi et al., 2022; Jumper et al., 2021). *Pymol v2.5.0 was used to render the structure*().

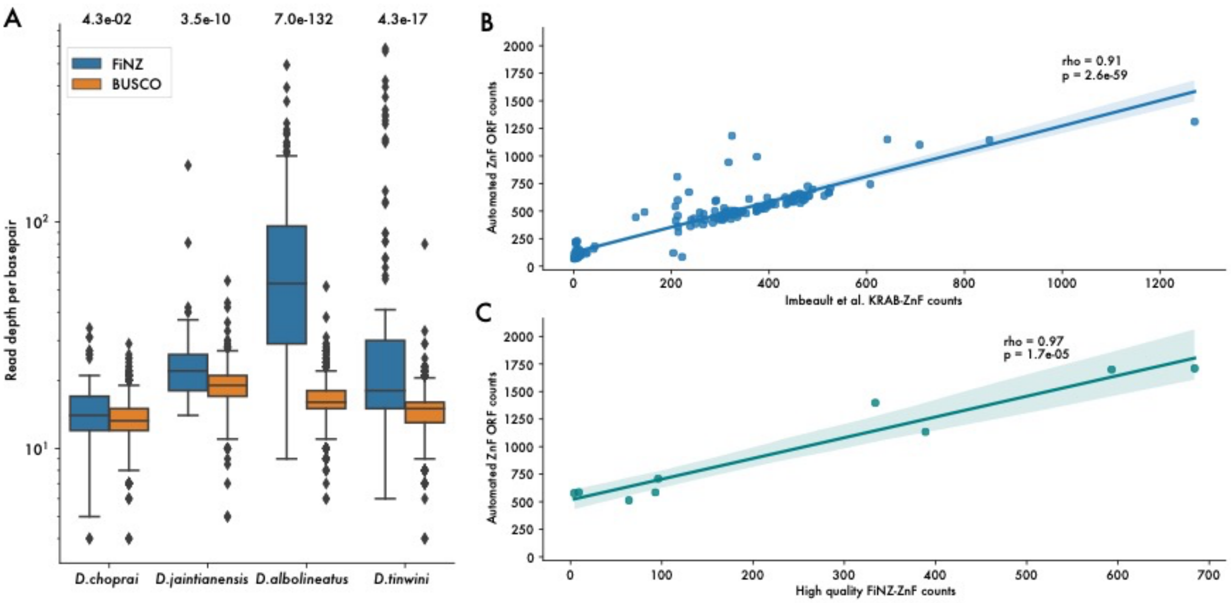


***Supplementary Figure 3. Quality control analyses for ZNF gene annotations.***

*A) Comparison of read depth over annotated BUSCO and annotated FZNF genes. Median read depth varies between species, but is generally higher over FZNF genes, and sometimes dramatically so, Danio albolineatus, read depth is approximately 50% higher than that of BUSCO genes, and therefore annotate gene counts should be treated as lower bounds on the true number. B) Spearman's rank correlation between automated ZNF ORF counts and independently generated counts from Imbeault et al.* (Imbeault et al., 2017). C) *Spearman's rank correlation between automated ZNF ORF counts and improved FZNF annotations. Taken together, these analyses indicate that genome assembly quality is likely to be the limiting factor in predicting ZNF genes, rather than annotation method.*

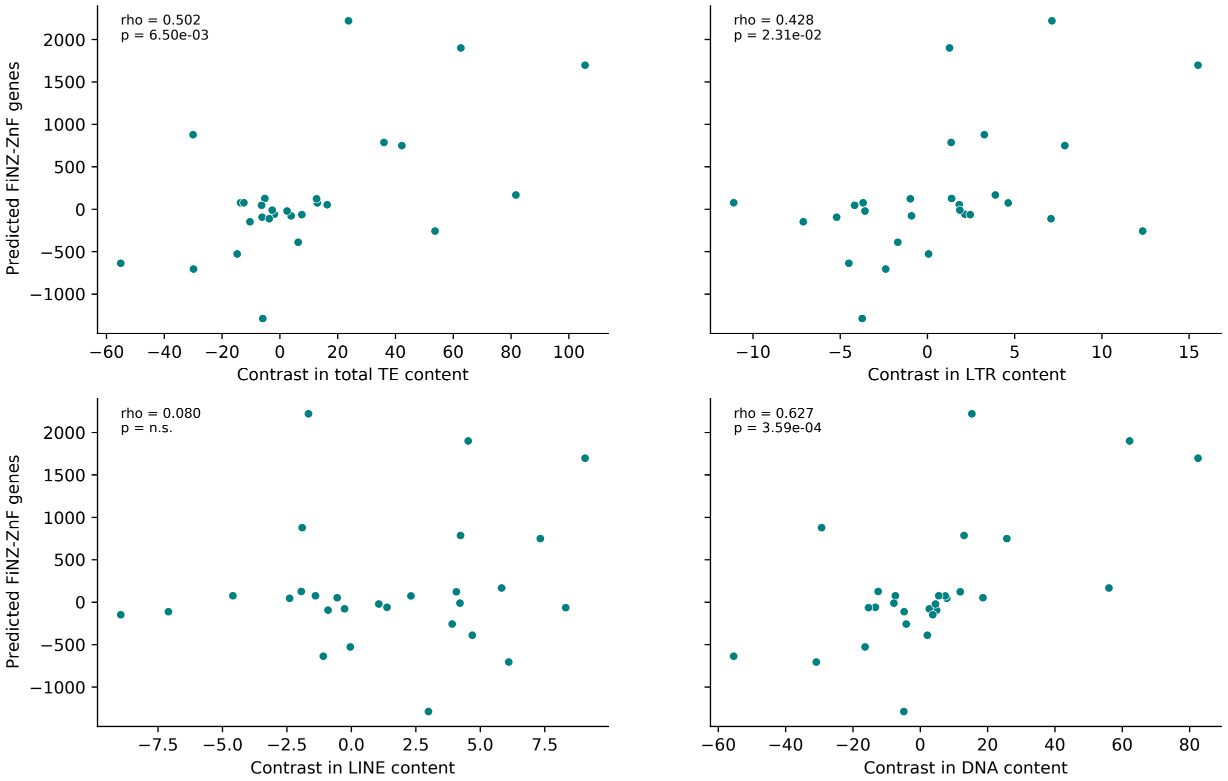


***Supplementary Figure 4. Phylogenetically independent contrasts between FZNF copy number and TE coverage***

*This figure relates to Fig. 2C and demonstrates that the correlation between FZNFs and TEs is not restricted to retroelements. Correlation calculated with Spearman's rank correlation test on phylogenetically independent contrasts between FZNF count and TE class genomic coverage.*

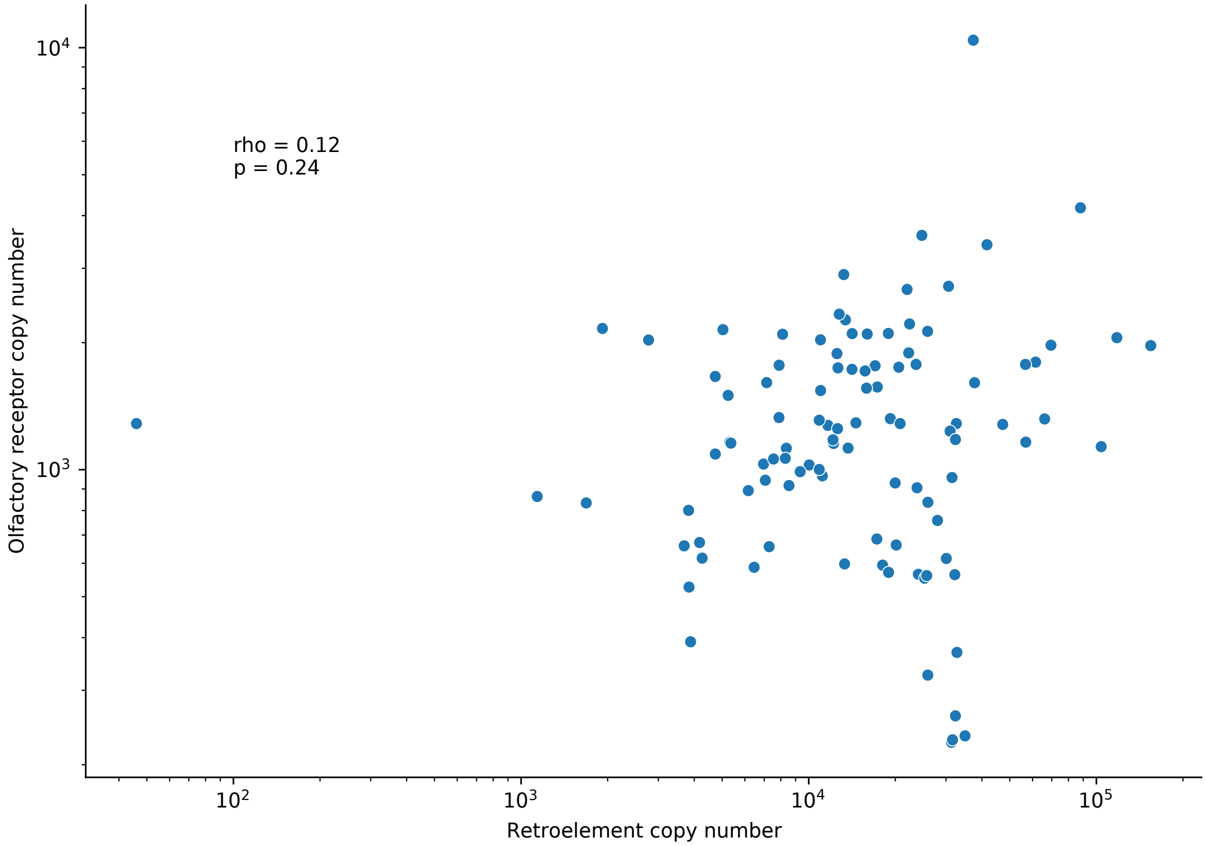


***Supplementary Figure 5. Correlation between retroelement and olfactory receptor copy number.*** *There is no significant correlation (Spearman's rank) between retroelement copy number and that of mammalian olfactory receptors, indicating that there is not a general relationship between TE content and gene copy number for large, multi-copy gene families. Each point is a representative species from a mammalian family.*

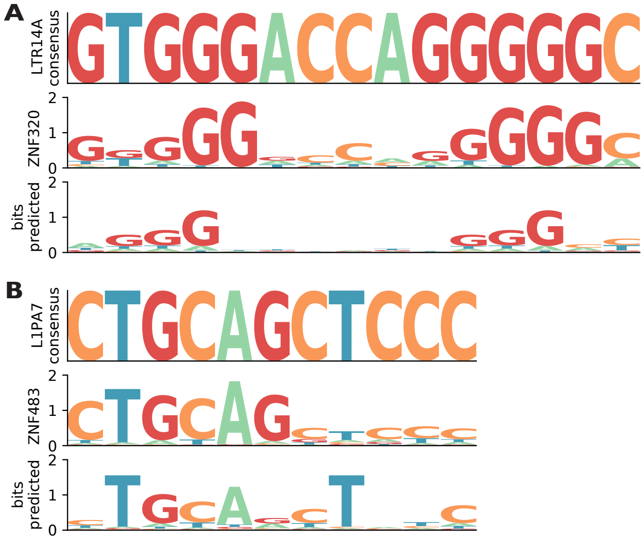


***Supplementary Figure 6. Examples of experimentally determined and predicted ZNF binding specificity to human TEs.*** *A) ZNF320 binding site on LTR14A consensus sequence. B) ZNF483 binding site on L1PA7. Y-axis records Shannon entropy (bits) for experimentally determined (2nd row) and predicted (3rd row) motifs.*

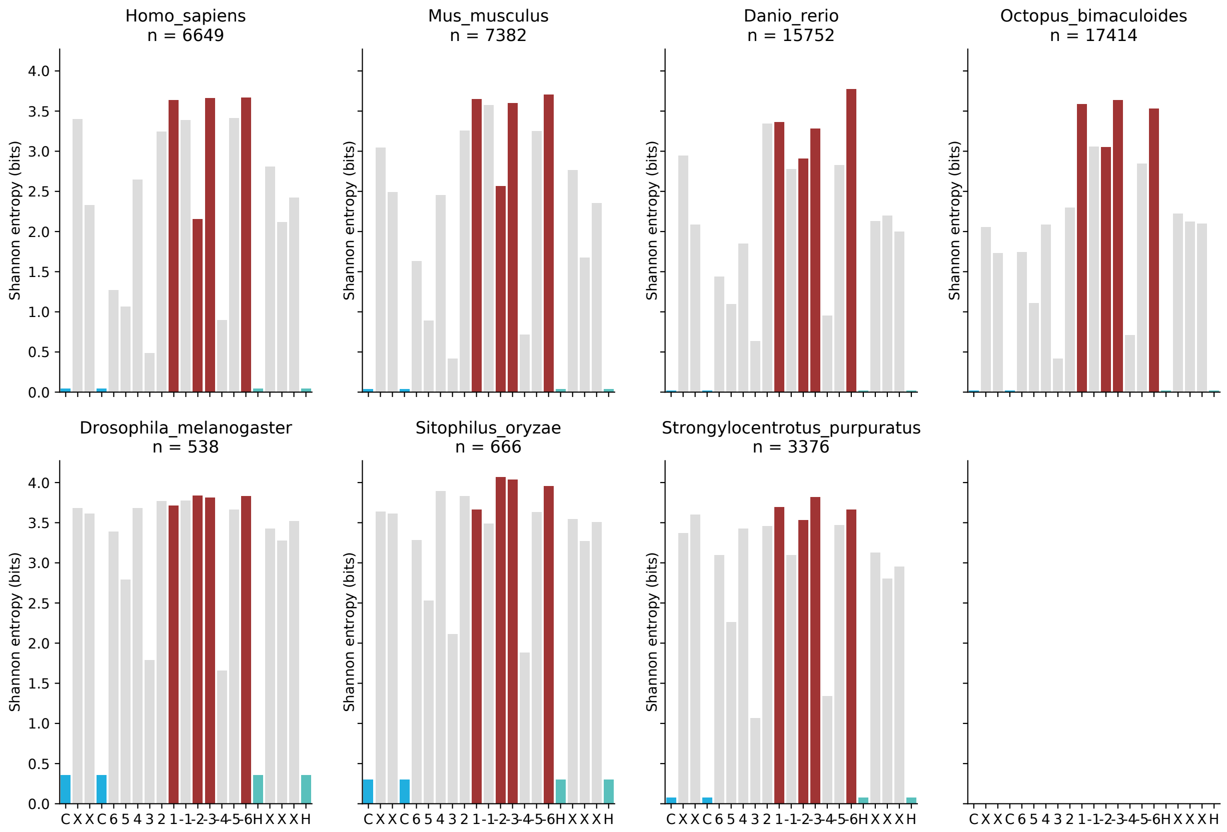


***Supplementary Figure 7. Sequence Shannon entropy at positions within the canonical ZNF domain.*** *Analysis of Shannon entropy reveals that sequence disorder (i.e. randomness) is highest at base-contacting residues within individual ZNF motifs in each species. This observation is consistent with rapid sequence turnover at these sites, in response to selection for varied nucleotide binding capabilities.*

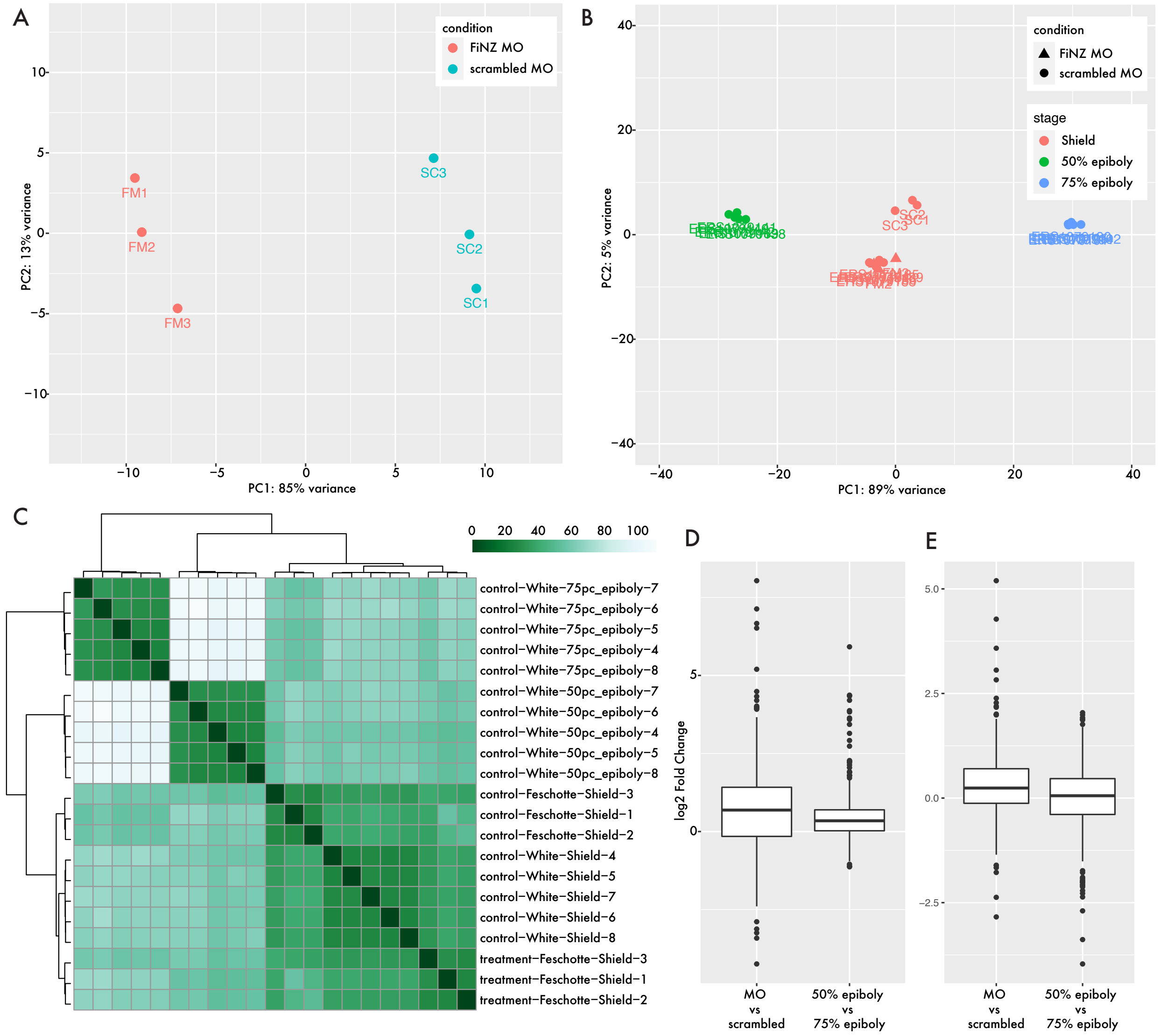


***Supplementary Figure 8. Comparison of treatment vs control and between-stage differential gene expression.***

*A) Principal component analysis showing separation of zebrafish embryo batches treated with FiNZ-targeting Morpholino or scrambled control Morpholinos. B) Principal component analysis using independently* (White et al., 2017) *collected zebrafish samples from 50% and 70% epiboly stages, demonstrating that the separation between FiNZ Morpholino and scrambled batches is not well explained by differences in embryo staging (i.e. the timing at which different batches were collected). C) Hierarchically clustered heatmap of distances between samples (after variance stabilizing transformation ). "White" and "Feschotte" refer to the lab of origin for each sample. D) Boxplot comparing log2 fold-change in expression of FZNFs for Morpholino vs. scrambled and 50% vs. 75% epiboly samples. E) As for D, but observing fold-change LTR family expression. D and E illustrate that the magnitude of changes in expression cannot be explained by stage alone.*

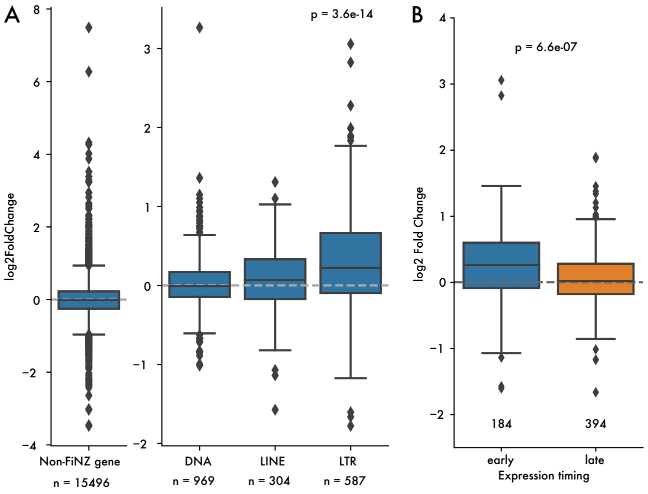


***Supplementary Figure 9. Comparison of differences between TE classes and expression stages***

*A) Log2 fold-change for all non-FiNZ genes, and comparisons between TE classes, demonstrating that LTRs are driving most of the signal for changes in TE expression. B) Log2 fold-change comparison between early and late-expressed TE families, demonstrating that only those families whose expression peaks prior to shield stage (where we collected embryos) have visible changes in expression.*
