## Supplementary Data 2 for "Transposable elements drive the evolution of metazoan zinc finger genes": README.md

This dataset contains predicted ZNF and retroelement ORFs, along with higher
quality gene annotations for cyprinid FiNZ-ZNFs. For zebrafish, which were
annotated separately, predicted 5' and 3' UTRs are included, and the RefSeq
genome assembly (GCF_000002035.6_GRCz11) was used, rather than the GenBank
assembly. Alternative haplotypes were removed from this file prior to
annotation.

`beds/` contains automated ORF predictions in BED file format
`gffs/` contains cyprinid Augustus FiNZ-ZnF annotations in GFF format
