## Supplementary Data 3 for "Transposable elements drive the evolution of metazoan zinc finger genes": README.md

This data file contains all the ZNF binding predictions for ZNF ORFs identified
as part of our high throughput annotation set (see BED files in Supp. Data 2).
The results of tests for binding motif enrichments are located in the `.tsv` 
files found in the `meme-out` directory.
