## Supplementary Data 4 for "Transposable elements drive the evolution of metazoan zinc finger genes": README.md

This data file contains the results of tests for positive selection acting on
Danio rerio FZNF paralogs. A neighbor-joining tree was generated from pairwise
distances for all pairs of cyprinid FZNFs, and seven zebrafish-specific clades
of in-paralogs were selected from this. These were realigned using PRANK,
trimmed to remove any gaps. Trees were generated using IQ-TREE, and these were
used as input for dN/dS analyses in PAML. Final results are given in 
`hypothesis_testing.xlsx`.
