## Supplementary Data 5 for "Transposable elements drive the evolution of metazoan zinc finger genes": README.md

# FiNZ-ZnF annotations and expression data

The gene annotations themselves are in the .gff file. You can safely ignore the
"interblock" or "protein match" lines, as these are just output from Augustus,
the tool I mostly used to generate the annotations. There should be 684 genes in
total, although this number is somewhat arbitrary since changing parameters will
lead to more or less being identified. Overall, many more genes are annotated in
this set than Ensembl or RefSeq, but that probably comes with a higher number of
non-functional/pseudogenes. I have also annotated UTRs, but these are tricky to
predict so I would be careful about relying on them too much.

For the expression data, I have remapped two datasets, the first being from
White et al., eLife (2017) and the second being Winata et al., Development (2018).
White et al has many more replicates and time points, but relies on
poly-A pull-down to capture reads, and so is less informative for maternally
deposited genes, hence why I looked at Winata et al, which focuses specifically
on maternally deposited genes.

For both datasets I used the same mapping approach: briefly, I used STAR for
mapping reads, allowing multi-mappers. Read-counting was done with
TEtranscripts, a tool from Molly Hammel's group. Calculation of TPM and other
downstream analysis was done in python, and I'm happy to share any other
details if that's useful to you! You should be able to get the different 
clusters for each gene in the "finz_expression_tpm.txt" file. Both files contain
TPM info for both genes and TEs, and FiNZ genes should be easily identifiable as
they are named "gXYZ". e.g. `grep "^g" finz_expression_tpm.txt` should get them
all.
